# Spousal Concordance in Telomere Length: New Evidence from Older Adults in the US

**DOI:** 10.1101/327528

**Authors:** Jason M. Fletcher

## Abstract

Telomere length (TL) has been associated with a range of aging outcomes as well as mortality. Recent research has shown both high heritability (∼70 %) of TL as well as moderate spousal similarity (r∼.3) using European datasets. This paper explores the level of spousal concordance in telomere length in the Health and Retirement Study, a national sample of adults in the US. The results both show that the spousal similarity in TL is lower in the US and also varies by the length of time spouses have been married as well as the educational attainments of husbands. These findings suggest the possibility of both assortative mating patterns related to telomere length as well as likelihood of shared environmental factors that cause similarity in TL in people who are socially connected.

## Background

Telomeres are repetitive DNA structures located at the ends of chromosomes that are thought to maintain genomic stability and shorten over time with each cell division (Hodes et al. 2002). They have been used as a measure of biological aging that has been shown in some (but not all) studies to be associated with mortality (Cawthon et al. 2003; Martin-Ruiz et al. 2005; Bischoff et al. 2006; Fitzpatrick et al. 2011). Some have proposed that they may be a marker of “healthy aging” rather than one of survival (Terry et al. 2008; Njajou et al., 2009), though either explanation may be related to the tendency for women to have greater TL than men (Barrett and Richardson, 2011; Honig et al., 2012; Shaffer et al., 2012; Zhu et al., 2011). There is growing interest in understanding the genetic factors, including the patterns of paternal vs. maternal inheritance, that predict telomere length (TL) as well as whether there are important environment and gene-environment interaction effects.

Some work has suggested the relatively large importance of biological factors as well as early environments that might shape TL over the life course. Broer et al. (2013) meta-analyze data from several cohorts to estimate a heritability of TL around 70% and also show evidence of stronger mother-child correlations in TL than father-child correlations. Hjelmborg et al. (2015) note that TL attrition is much slower in adults than in children and that having a long or a short TL is largely determined before adulthood. The authors find that heritability and early life environment are the main determinants of TL throughout the human life course.

While the analysis in Broer et al. (2013) focuses on genetic pathways, a secondary set of results focused on spousal similarity in TL and found moderate correlations (r∼.3). This estimate is the first in the literature and has not been extended by other researchers. Broer et al. (2013) suggest a few possible interpretations of this correlation. First, the resemblance could be induced by living together for a long time. A second possibility is an ascertainment effect—focusing on older adults who are married (and thus survived into old age) will produce a sample who is likely to have above average TL. A third possibility, not raised by Broer et al, is of genetic assortative mating. Domingue et al. (2014) suggest a moderate level of correlation across the genomes of spouses, compared with unrelated non-spouses.

Depending on the pathways that produce spousal similarity in TL, assessments of heritability of TL as well as intergenerational analysis become more difficult. Non-random mating patterns of parents can lead to heritability estimates that are understated. These assortative mating patterns may also make estimated associations between paternal-child and maternal-child TL difficult to interpret. In contrast, TL spousal similarity may reflect a largely environmental pathway, which could suggest a larger set of interventions during older adulthood that could benefit longevity than is suggested by Hjelmborg et al and other researchers as well as be used as a clinical signal to investigate the health of spouses (and possibly other family members and neighbors or co-workers) of individuals with low TL.

In order to explore these possibilities, this paper conducts the second investigation of spousal associations in TL in the literature and the first investigation with US data. Using the Health and Retirement Study, this paper analyses data on nearly 1,500 spousal pairs (Broer et al. use over 1,900 spouses across 5 datasets) and finds spousal correlations in TL that vary by spousal characteristics. In particular, spousal TL correlations are higher for couples who have one marriage than those involved in their second (or higher) marriage. This suggests assortative mating may play a smaller role than shared environment in producing spousal TL correlations.

### Data

The Health and Retirement Study began in 1992 as a nationally representative longitudinal study of aging of individuals born 1931-1941 and their spouses (Sonnega et al. 2014). The focus of the first and subsequent surveys has been in collecting data on the aging process by focusing on health, work, family, and related domains. There are now >10,000 individual with genetic data collected. Of these, nearly 6,000 have telomere data, where the subjects were selected based on membership in the one-half random sample of the HRS that were preselected to have enhanced face-to-face interviews in 2008.^2^ (see Faul et al. 2016 for additional details on assessment genotyping).

Descriptive statistics for the sample of spouses in the HRS who have telomere assessments are presented in Table 1, which is stratified by gender. Wives in the sample are on average 65.5 years old and completed 12.7 years of schooling. Husbands are 68.7 years old on average and have completed 12.9 years of schooling. Nearly 2/3rds of the respondents report being married only once. Appendix Table 1 shows that the full sample of respondents are quite similar to the sample of spouses in TL, age, and educational attainment and also shows the descriptive statistics of the pooled spousal sample.

**Table 1.** Descriptive Statistics of Spouses Characteristics and Telomere Length in the HRS

Wives (N~1500)
| Variable | Mean | Std Dev | Min | Max |
| --- | --- | --- | --- | --- |
| Telomere Length | 1.35 | 0.32 | 0.23 | 2.96 |
| Age (2008) | 65.49 | 9.78 | 33.00 | 90.00 |
| Educational Attainment | 12.71 | 2.96 | 0.00 | 17.00 |
| White | 0.86 | 0.35 | 0.00 | 1.00 |
| Black | 0.08 | 0.28 | 0.00 | 1.00 |
| Other Race | 0.06 | 0.23 | 0.00 | 1.00 |
| Number of Marriages | 1.44 | 0.76 | 0.00 | 7.00 |
| Single Marriage | 0.67 | 0.47 | 0.00 | 1.00 |

Husbands (N~1500)
| Variable | Mean | Std Dev | Min | Max |
| --- | --- | --- | --- | --- |
| Telomere Length | 1.29 | 0.33 | 0.23 | 2.85 |
| Age (2008) | 68.69 | 9.43 | 26.00 | 93.00 |
| Educational Attainment | 12.88 | 3.36 | 0.00 | 17.00 |
| White | 0.86 | 0.35 | 0.00 | 1.00 |
| Black | 0.09 | 0.28 | 0.00 | 1.00 |
| Other Race | 0.05 | 0.22 | 0.00 | 1.00 |
| Number of Marriages | 1.46 | 0.75 | 0.00 | 7.00 |
| Single Marriage | 0.65 | 0.48 | 0.00 | 1.00 |

Notes: observations with Telomere Length above 3 are not included.

## Results

Table 2 presents the key empirical results that focus on spousal similarity in TL. The adjusted correlation between spouses TL is 0.085, which is smaller than the reports in Broer et al. (2013) of ∼0.3 using European samples. As expected, Husband age is negatively related to husband TL; otherwise, few wife or husband characteristics predict TL (see Appendix Table 2A for additional analysis of individual level predictors of TL). In order to explore alternative hypotheses for the sources of the correlations in spousal TL, columns 2 and 3 stratify the sample based on whether the husbands have been married once or more than once. The results show much stronger adjusted correlations in spousal TL among those spouses married once (0.12) than those married more than once (0.026), this difference is statistically significant at the 9% level.

**Table 2.** Adjusted Associations between Spousal Telomere Length in the HRS Full Sample and Stratified by Husband’s Number of Marriages

| Outcome | TL | TL | TL |
| --- | --- | --- | --- |
| Sample | Spousal | Husband Single Marriage | Husband 1+ Marriages |
| Spousal TL | 0.085***<br>(0.027) | 0.119***<br>(0.034) | 0.026<br>(0.043) |
| Husband Education | -0.001<br>(0.003) | 0.002<br>(0.004) | -0.009*<br>(0.005) |
| Husband Black Race | 0.094<br>(0.082) | 0.038<br>(0.105) | 0.194<br>(0.133) |
| Husband Other Race | 0.042<br>(0.045) | 0.025<br>(0.064) | 0.081<br>(0.066) |
| Husband Single Marriage | 0.020<br>(0.025) |  |  |
| Wife Education | 0.005<br>(0.004) | 0.003<br>(0.004) | 0.009<br>(0.006) |
| Wife Black Race | -0.006<br>(0.084) | 0.076<br>(0.108) | -0.153<br>(0.135) |
| Wife Other Race | 0.006<br>(0.043) | 0.025<br>(0.063) | -0.016<br>(0.059) |
| Wife Single Marriage | -0.005<br>(0.025) | 0.005<br>(0.038) | 0.003<br>(0.033) |
| Husband Age | -0.004**<br>(0.002) | -0.004<br>(0.003) | -0.003<br>(0.002) |
| Wife Age | -0.000<br>(0.002) | -0.001<br>(0.003) | 0.000<br>(0.002) |
| Constant | 1.414***<br>(0.091) | 1.398***<br>(0.119) | 1.463***<br>(0.147) |
| Observations | 1,492 | 975 | 517 |
| R-squared | 0.038 | 0.050 | 0.028 |
| Chi2 |  |  | 3.042 |
| P |  |  | 0.081 |

Notes: each observation is a spousal pair. Standard errors in parentheses
\*\*\* p<0.01, \*\* p<0.05, \* p<0.1

Table 3 further stratifies the results by educational attainment of the spouse. For husbands with low education (column 1), the spousal correlations are lower (0.05) than husbands with higher education (0.12 in column 2), though the p-value for this differences is p<0.18. In contrast there is no evidence of differences in spousal similarity in TL that varies by wife’s educational attainment (columns 3 and 4).

**Table 3.** Adjusted Associations between Spousal Telomere Length in the HRS Stratified by Education Level

| Outcome | TL<br>Husband<br>HS grad<br>or Less | TL<br>Husband<br>Greater than HS<br>Grad | TL<br>Wife<br>HS grad or<br>Less | TL<br>Wife<br>Greater than HS<br>Grad |
| --- | --- | --- | --- | --- |
| Sample |  |  |  |  |
| Spousal TL | 0.049<br>(0.038) | 0.124***<br>(0.038) | 0.074**<br>(0.033) | 0.087**<br>(0.039) |
| Husband Education | -0.009*<br>(0.005) | -0.007<br>(0.009) | 0.002<br>(0.004) | 0.001<br>(0.005) |
| Husband Black Race | 0.146<br>(0.121) | 0.034<br>(0.112) | 0.033<br>(0.113) | -0.033<br>(0.114) |
| Husband Other Race | 0.102<br>(0.065) | -0.032<br>(0.062) | -0.016<br>(0.056) | 0.007<br>(0.070) |
| Husband Single Marriage | 0.002<br>(0.036) | 0.034<br>(0.035) | 0.084**<br>(0.033) | 0.015<br>(0.036) |
| Wife Education | 0.006<br>(0.005) | 0.004<br>(0.005) | -0.007<br>(0.005) | -0.006<br>(0.009) |
| Wife Black Race | -0.053<br>(0.121) | 0.051<br>(0.121) | 0.057<br>(0.116) | 0.158<br>(0.115) |
| Wife Other Race | -0.050<br>(0.060) | 0.051<br>(0.063) | 0.024<br>(0.055) | 0.045<br>(0.065) |
| Wife Single Marriage | -0.015<br>(0.036) | 0.011<br>(0.035) | -0.047<br>(0.032) | -0.008<br>(0.037) |
| Husband Age | -0.006**<br>(0.002) | -0.001<br>(0.002) | 0.001<br>(0.002) | 0.003<br>(0.003) |
| Wife Age | 0.000<br>(0.002) | -0.002<br>(0.002) | -0.006***<br>(0.002) | -0.009***<br>(0.003) |
| Constant | 1.612***<br>(0.134) | 1.366***<br>(0.164) | 1.592***<br>(0.117) | 1.656***<br>(0.171) |
| Observations | 751 | 741 | 820 | 672 |
| R-squared | 0.049 | 0.039 | 0.047 | 0.062 |
| Chi2 |  | 1.876 |  | 0.060 |
| P |  | 0.171 |  | 0.806 |

Notes: each observation is a spousal pair. Standard errors in parentheses
\*\*\* p<0.01, \*\* p<0.05, \* p<0.1

## Conclusions

TL has been shown to be an important predictor of old age health and potentially linked to mortality. However, the key determinants of TL are largely unknown. On one hand, heritability estimates of TL are often in the range of 70%, suggesting the likelihood of important genetic factors. Related evidence has shown that early childhood conditions are strong predictors of late life TL. These findings could suggest that late life interventions to reduce TL shortening may be ineffective. On the other hand, spousal similarity in TL (of unrelated people) may instead suggest environmental factors continue to shape TL in middle and older age individuals and point the possibility of effective interventions during the middle and later stages of the life course. The findings in this study present new evidence of the possible importance of these later life factors. Using a large set of spouses from a national sample of older individuals in the US, this study shows modest correlations in spousal TL as well as evidence that couples who have been together longer have higher levels of similarity in their TL. Findings also suggest that spousal similarity in TL is lower in couples with less educated husbands. Additional research is needed that explores potential determinants of these environmentally induced TL patterns.

## Appendix Tables

**Appendix Table 1.** Descriptive Statistics of Full Sample and Pooled Spousal Sample

| Variable | Obs | Mean | Std Dev | Min | Max |
| --- | --- | --- | --- | --- | --- |
| Telomere Length | 5733 | 1.32 | 0.33 | 0.20 | 2.98 |
| Female Indicator | 5733 | 0.59 | 0.49 | 0.00 | 1.00 |
| Age (2008) | 5733 | 68.77 | 10.34 | 26.00 | 107.00 |
| Educational Attainment | 5725 | 12.54 | 3.19 | 0.00 | 17.00 |
| White | 5733 | 0.82 | 0.39 | 0.00 | 1.00 |
| Black | 5733 | 0.13 | 0.34 | 0.00 | 1.00 |
| Other Race | 5733 | 0.05 | 0.22 | 0.00 | 1.00 |
| Number of Marriages | 5721 | 1.40 | 0.77 | 0.00 | 7.00 |
| Single Marriage | 5733 | 0.65 | 0.48 | 0.00 | 1.00 |
| Telomere Length | 3022 | 1.32 | 0.33 | 0.23 | 2.96 |
| Female Indicator | 3022 | 0.50 | 0.50 | 0.00 | 1.00 |
| Age (2008) | 3022 | 67.09 | 9.74 | 26.00 | 93.00 |
| Educational Attainment | 3019 | 12.79 | 3.17 | 0.00 | 17.00 |
| White | 3022 | 0.86 | 0.35 | 0.00 | 1.00 |
| Black | 3022 | 0.09 | 0.28 | 0.00 | 1.00 |
| Other Race | 3022 | 0.05 | 0.23 | 0.00 | 1.00 |
| Number of Marriages | 3021 | 1.45 | 0.76 | 0.00 | 7.00 |
| Single Marriage | 3022 | 0.66 | 0.47 | 0.00 | 1.00 |

Notes: observations with Telomere Length above 3 are not included.

**Appendix Table 2.** Associations between Individual Characteristics and Telomere Length in the HRS

| Sample | TL<br>Full | TL<br>Spousal | TL<br>Wives | TL<br>Husbands |
| --- | --- | --- | --- | --- |
| Female Indicator | 0.028***<br>(0.009) | 0.042***<br>(0.012) |  |  |
| Age (2008) | -0.005***<br>(0.000) | -0.005***<br>(0.001) | -0.006***<br>(0.001) | -0.005***<br>(0.001) |
| Educational Attainment | 0.001<br>(0.001) | 0.001<br>(0.002) | 0.001<br>(0.003) | 0.001<br>(0.003) |
| Black Race | 0.094***<br>(0.013) | 0.108***<br>(0.021) | 0.113***<br>(0.030) | 0.104***<br>(0.030) |
| Other Race | 0.023<br>(0.020) | 0.040<br>(0.026) | 0.034<br>(0.036) | 0.047<br>(0.039) |
| Number of Marriages | -0.019***<br>(0.006) | -0.008<br>(0.008) | -0.009<br>(0.011) | -0.007<br>(0.011) |
| Constant | 1.631***<br>(0.039) | 1.645***<br>(0.053) | 1.725***<br>(0.073) | 1.604***<br>(0.075) |
| Observations | 5,715 | 3,019 | 1,508 | 1,508 |
| R-squared | 0.038 | 0.044 | 0.044 | 0.030 |

Standard errors in parentheses. \*\*\* p<0.01, \*\* p<0.05, \* p<0.1

## Footnotes

1 The Health and Retirement Study (HRS accession number 0925-0670) is sponsored by the National Institute on Aging (grant numbers NIA U01AG009740, RC2AG036495, and RC4AG039029) and is conducted by the University of Michigan. Additional funding support for genotyping and analysis were provided by NIH/NICHD R01 HD060726. I thank Angela Forgues for excellent research assistance.

2 Faul et al. (2016) note that this sample was selected at the household level to ensure that the same request was made to both members of a household. There were a small number of exclusion criteria: (1) needing to be interviewed by proxy, (2) residing in a nursing home, or (3) preferring to be interviewed by telephone. Respondents who provided a saliva sample did not differ by age, sex, education, or income from those who were asked but did not consent (Faul et al. 2016). Consent for the DNA collection was obtained in person at the time of the interview. The HRS study protocol was approved by the University of Michigan Institutional Review Board.

